# Using Natural Vector Method for Population Genomic Analysis on Human Mitochondrial Genome Data

**DOI:** 10.64898/2026.07.11.737899

**Authors:** Mengcen Guan, Qi Wu, Xin Zhao, Stephen S.-T. Yau

## Abstract

The natural vector method is an important method for the analysis of biological sequences. In this study, we applied this method to population genetic analysis, with the core purpose of using it to evaluate the characteristics of a set of sequences rather than just pairwise comparison. We used the mitochondrial genome dataset from the human 1000 Genomes Project as a dataset to verify the feasibility of this improved natural vector method. The results showed that the modified natural vector method could be used for various population genetic approaches at least in the sense of population average, including the calculation of principal component analysis, population structure analysis and genetic diversity parameters. The results were in good agreement with those based on traditional molecular genetic markers such as SNP. The new method validates the feasibility of natural vector method for population genetic analysis and provides a framework for the application of matchless pair method to population genomic analysis on a wider scale.

## 1 INTRODUCTION

In evolutionary biology, macroevolution studies the evolution of taxa above species. At the molecular level, the phylogenetic tree is usually used to represent the phylogenetic relationship between sequences in a sample taken from a higher-order taxon. On the contrary, microevolution is the study of evolution at the population level below the species level. At the molecular level, for a sequence sample including multiple individuals, a variety of methods can be used to quantify the sequence divergence or discripancy to characterize the genetic diversity and structure of the population from which the sample comes [1].

Alignment-free methods constitute a broad category of approaches for sequence analysis that do not require alignment between sequences [2, 3]. These methods extract subsequences from the sequences and provide a quantification of the sequences based on the numerical features of these subsequences, followed by sequence analysis based on these quantification. Here the “subsequences” typically refer to individual nucleotides or oligonucleotide kmers, while the “numerical features” commonly used are the frequencies or coordinates of these subsequences within the sequences. The resulting quantification often takes the form of a high-dimensional vector. Over the past decades, alignment-free sequence analysis methods have developed rapidly, demonstrating strong capabilities in capturing global sequence characteristics and enabling rapid sequence classification. Although such methods have been widely used in genome phylogenetic analysis, metagenomic analysis and so force, few works have applied alignment-freee methods for population level sequence analysis. This is because the main output of alignment-free methods is the construction of distance trees between sequences, while population genomic analyses, as mentioned above, often requires more detailed description or representation of the information in a sample of sequences, rather than just using trees to show the relationship between sequences in a sample.

The Natural Vector method [4]is an approach to alignment-free sequence analysis, distinguished by its use of the coordinates of the four nucleotides within a sequence as numerical features, as well as the construction of numerical vectors using the n-th order moments of these coordinates. For a given sequence, the counts of the four nucleotides, along with the four means (first-order moment) and the four variances (second-order moment) of their coordinates, constitute a 12-dimensional vector describing the sequence. When considering more detailed scenarios, each additional order moment increases the vector’s dimensionality by four. This method demonstrates computational speed advantages in analyses ranging from DNA sequences to protein structures. Like the other alignment-free methods mentioned above, natural vector method mainly focuses on the pairwise difference comparison when applied to a set of sequence samples for sequence analysis. The difference relationship between sequences is described by using a tree structure. The limitation of this in application is that it can only be used in the field of phylogenetic-like analysis and is difficult to carry out population genetic analysis.

In this paper, we attemped to apply the natural vector method to population-level genomic analyses. By analogy with the “location-allele” theoretical framework of genetics, the dimension of the space where the numerical vector is located in the natural vector method is compared to “loci”, and the numerical segments in the dimension are compared to “alleles”. In this way, the sequence data after the change of the natural vector method is compared to a group of multi-allelic genetic loci. We used such an idea to perform population genomic analysis on data from the human mitochondrial genome dataset from the 1000 Genomes Project. We did three analyses, principal components, population structure, and genetic diversity calculations. The results are compared with those of classical SNP-based analyses. The results showed a good parallel between the results obtained by the natural vector method and the SNP-based results. Our approach extends the application of natural vector method to real biological data and provides a new potential analysis method for population genomics data.

## 2 MATERIALS AND METHODS

### 2.1 Natural Vector

The natural vector is a 12-dimensional numerical vector used to encode a DNA sequence and describe the distribution of the nucleotides A, G, C, and T[4]. The definition is as follows. Given a sequence with a length of n: *S* = *s*_1_*s*_2_ *. . . s_n_ , s_i_* ∈ {*A, G, C, T* }, the indicator functions for A, C, G, T (*w_k_*(.)*, k* ∈ {*A, G, C, T* }) are defined as:

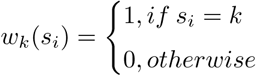

The indication functions describe the position information of the four nucleotides in sequences. The components of the natural vector can be calculated based on these functions:

- The counts of nucleotides, denoted as *n_k_*, are given by 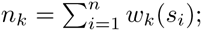
- The average locations, denoted as *µ_k_*, are given by 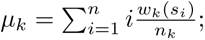
- The second central moment of positions, denoted as 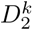, are determined by 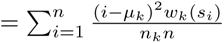.

The 12-dimensional natural vector is composed by these three components:

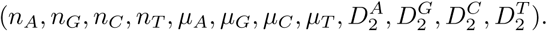

The counts, average locations and the central moments of four nucleotides A, G, C, T are natural parameters associated to a DNA sequence. And these combined numerical parameters are sufficient to characterize each DNA sequence, so these parameters give a complete understanding of four nucleotides A, G, C and T.

### 2.2 Embodying a generalized “locus-allele” Paradigm in natrual vector methods

Assuming a population genomics dataset with N individuals from M predefined geographic or phenotypic populations, converting each sequence into a natural vector results in N×12 data, where 12 is the vector’s dimension. These dimensions can be analogized to genetic loci, establishing the concept of generalized loci (”locus”). Thus, all sequences in a sequence sample can be quantitatively analyzed in one single dimension. In the example with N individuals, there are 12 dimensions. The natural vector’s dimensions, designed as multiples of 4 (e.g. 16, 20), yield correspondingly larger dimension space (N×16, N×20, etc.), and analyses can be performed in every single dimension.

One can study a sample of sequences by exploring the overall situation of the sample by some numerical characteristics of the sample. Therefore, different values of sequences in a natural vector dimension in a sample can be likened to different “allele states” in a “genetic locus” to explore its diversity. The first four dimensions represent nucleotide counts and are discrete integers, leading to a finite number of “alleles” for real sequences. For human mitochondrial genome data, these “alleles” states typically number around 10–100, matching the biological expectation for multiallelic loci. However, other dimensions involve real numbers, posing a challenge for defining allele frequencies, a central concept in population genetics. A solution is to partition these dimensions into intervals, treating sequences within the same interval as identical, thus defining an “allele”. The key then becomes devising a sensible interval partitioning strategy.

Population structure is common in most species, including subspecies distinctions and subdivisions within genetic populations. Often, populations defined by geography or phenotype are known in advance. Given this, we assume a clear population structure in the sample and use the ability to statistically identify it as a criterion for determining the optimal “allele” partitioning. For human mitochondrial genome data, we present a discretization algorithm as following, tailored to human population complexity, especially considering subdivisions within major racial categories. Since population structure is widespread, this strategy of partitioning “allele” to identify it is broadly applicable.

### 2.3 Discretization Algorithm

To transfer the concepts from traditional genetics to our new model, we need to generalize the concepts of “allele” and “locus.” In traditional genetics, a locus refers to one specific position of a gene on a chromosome, and an allele is one of the different forms of a gene at the same locus. The traditional loci and alleles of genetics are shown in Fig. S1 of supplementary material.

In the natural vector method, we define a “locus” as each position/dimension of the 12-dimensional natural vector. This is a natural generalization, so the generalized locus spans from the first to the twelfth dimension of the natural vector. Taking the number of adenine nucleotides (A) in a sequence as an example, since the count of this number occupying an unique dimension in the natural vector, we consider it (the “A” in the sequence) as one generalized “ locus” in the sequence.

To define an “allele” for the generalized locus in natural vector, two fundamental considerations should be necessarily addressed. First, within classical genetic theory, alleles are operationally defined in reference to specific chromosomal position, wherein their informational content strictly corresponds to localized genetic variation at designated loci. This locus-specific attribution must be preserved when extending the concept to generalized alleles. Second, regarding natural vector methods, the genetic diversity of a sample sequence from (a) population, represents as the diversity of statistical features of the sequences, and then represents as the observed numerical distribution of the dimensions of the natural vector. Consequently, generalized alleles must be operationally linked to:

- Discrete value states within natural vector dimensions (constituting generalized loci)
- Their corresponding frequency distributions across populations.

Suppose two sequences have the same numerical value at a dimension of natural vector, they are considered to belong to the same allele of the generalized locus. For instance, if two sequences A and B have the same count of adenine nucleotides, they share the same allele at that locus.

Although a natural definition method is that different integer values correspond to different generalized alleles, one can “arbitrarily” defined similar numerical values as the same generalized allele. Actually the definition of the natural vector have suggested that similar numerical values in a dimension indicate similar characteristics between sequences. As for integer values, this division of numerical intervals may be unnecessary, but for continuous real values, this division is necessary. That is the case of the last eight dimensions of the 12-dimensional natural vector, whose values are real number. In probability theory, when defining the probability of a continuous random variable, it is also necessary to consider an interval rather than defining the probability for a specific real number value. Assuming that each real value is treated as a unique allele, the number of alleles will become very large, and the frequency of each allele will be equal to the reciprocal of the population size, which is not meaningful in genetics. For this aim, we propose discretizing the continuous values of these eight dimensions. As shown in the Fig. S1, we can divide numerical values into equal intervals, using the midpoint of each interval to represent all values within it. These intervals then serve as the generalized alleles for the dimension.

So, the key issue is how to determine the number of discretized intervals in a natural way to avoid the impact of artificial partitioning. As mentioned in section 3.1, the considered standard is that the population has a known clear population genetic structure and the ability to statistically identify this population genetic structure is used as a criterion for determining the optimal allele discretization method. Specifically, we perform pairwise Wilcoxon rank-sum tests [5] between populations and select the number of intervals *n* that maximizes the differentiation between populations. Based on this criterion, we have designed an algorithm for discretizing the last eight dimensions of the 12-dimensional natural vector.

#### Algorithm 1 Discretization Algorithm

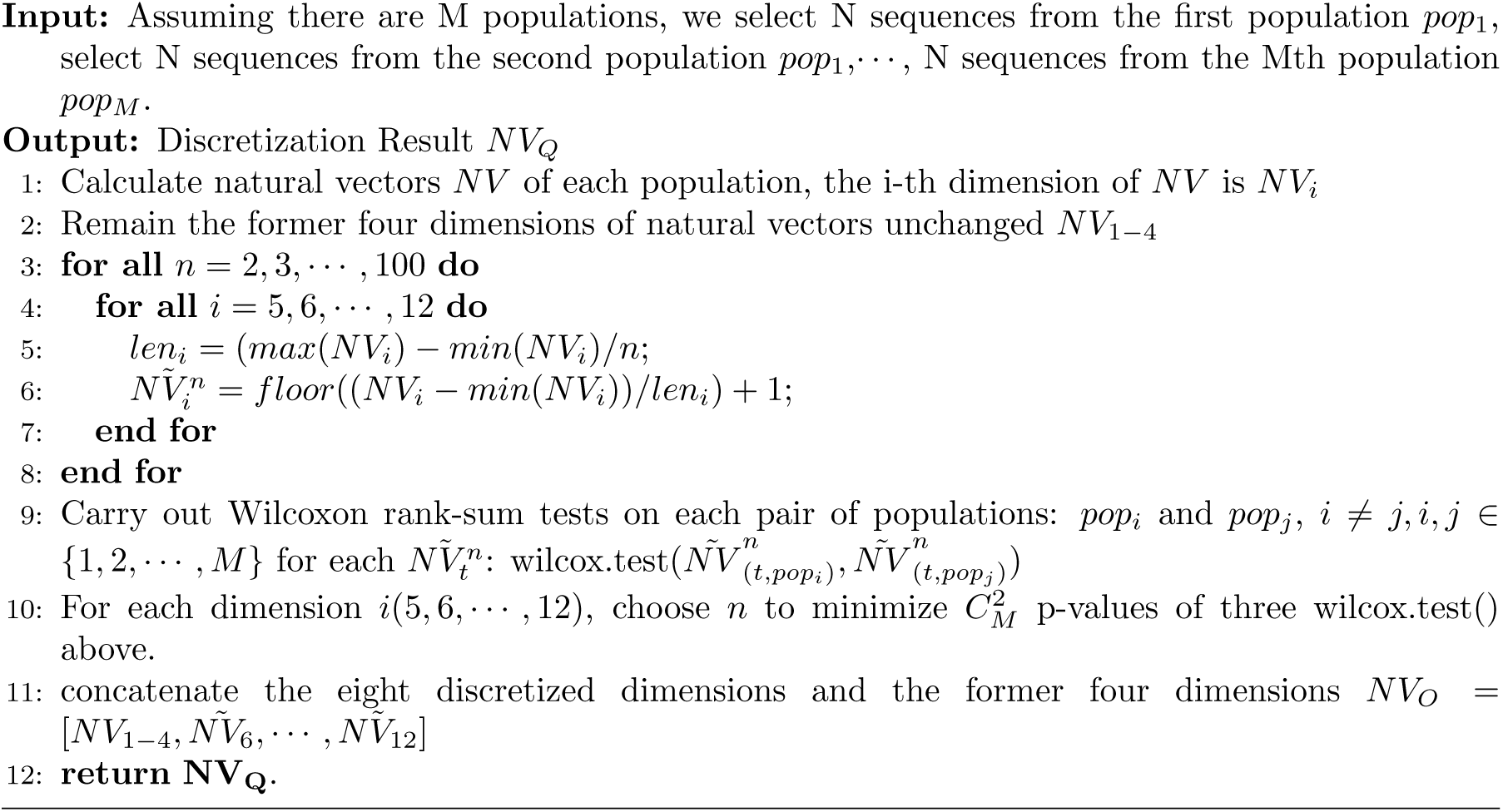

With such an approach we define a model with the generalized locus and the generalized allele as shown in sub-figure B of Fig. S1. We then firstly take three populations: British, Yoruba and Han Chinese as an example to show how we choose the number of intervals to discretize natural vectors. The core concept when designing the discretization algorithm is to furthest differentiate samples from different continents such as Asia and Africa.

### 2.4 The principal component analysis

To investigate the genetic structure among populations, we performed Principal Component Analysis (PCA) based on both mtDNA single nucleotide polymorphisms (SNPs) and natural vectors of mtDNA.

#### 2.4.1 PCA of mtDNA derived from natural vectors

Natural vectors encompass rich information from the original mitochondrial sequences and are one-to-one correspondence with the original sequences, genetic structure could be discovered through the PCA results of natural vectors of mtDNAs.

Natural vectors (12-dimensional) were derived from the mtDNA genomes of 2,534 individuals. Each vector was annotated with a population label corresponding to the sample’s origin. These vectors are not directly served as input for downstream PCA. Then mean vector of each population is calculated according to the definition 1.

##### Definition 1

**(mean vector)** *Given a collection of n vectors* 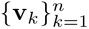 *in* R*^m^*, the ***mean vector*** *µ* ∈ R*^m^ is defined component-wise as:*

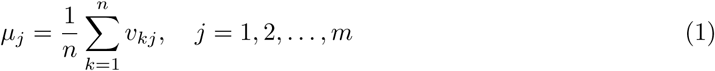

*µ_j_is the j-th element of the vector *μ*, *v_kj_* is the j-th element of the vector v_k_. or equivalently in vector form:*

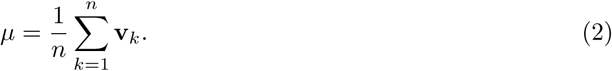

Then the mean vector of one population represents sequences from the population, all these mean vectors are input for PCA.

#### 2.4.2 PCA of mtDNA derived from SNP data

Mitochondrial SNP data were extracted from phased VCF files (1000 Genomes Project Phase 3). Processing of SNP data followed standard protocols.

PCA analysis of SNPs on population-level was performed by computing mean genotype vectors for each population. For population *k* with *n_k_*individuals, the population mean vector ***µ****_k_* was calculated as: where *S_k_* is the set of samples belonging to population *k*, and **v***_i_* is the genotype vector of the *i*-th individual, then the *µ_k_* is the mean vector defined in definition 1. The final population-average genotype matrix (**M** ∈ R*^p×m^* with *p* populations and *m* SNPs) was subjected to PCA.

### 2.5 The population structure analysis based on natural vectors

In the population structure analysis based on the natural vector method, the following procedure was adopted. First, for each population under study, the natural vectors of all individual sequences within that population were computed. Subsequently, the mean natural vector for each population was calculated to represent its central genetic characteristics.

Since the input for the STRUCTURE software requires discrete, integer values, the discretization algorithm was applied. This transformation converted the continuous natural vectors into a discrete, integer-based format compatible with STRUCTURE’s input requirements.

Finally, these discretized natural vectors, representing the genetic profiles of the different populations, were used as the input data for the STRUCTURE analysis. The software was then run to estimate the number of ancestral populations (K) and assign individual sequences or populations to these inferred genetic clusters, thereby visualizing and quantifying the underlying population stratification.

### 2.6 Measurement of Genetic Diversity based on natural vectors

Genetic diversity is an important component of biodiversity, which refers to the diversity of genetic characteristics in a species’ genetic composition, including genetic variation among different populations within a species or among different individuals within the same population. Here we consider the level of genetic diversity for different populations. Genetic diversity is the foundation of biodiversity and plays a crucial role in the survival, reproduction, and fitness of species to environmental changes [6, 7].

There are several methods for calculating the genetic diversity.

#### 2.6.1 Expected Heterozygosity

Gene diversity, also known as Expected Heterozygosity *H_e_*, refers to the probability that two alleles randomly drawn from one population are different. The calculation formula is as follows:

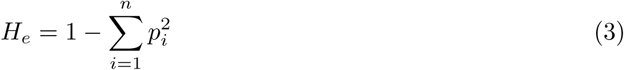

*p_i_*is the frequency of the i-th allele and *n* is the number of alleles. When a population is in Hardy-Weinberg equilibrium, genetic diversity is equal to expected heterozygosity.

#### 2.6.2 Average pairwise nucleotide diversity ***π***

Nucleotide diversity, denoted as *π*, is an important concept in molecular genetics and can serve as a crucial indicator of population genetics to measure the level of genetic diversity within a population. It can be calculated through definition 2.

##### Definition 2

(Average pairwise nucleotide diversity *π*) *Nucleotide diversity π represents the base differences per site on multiple sample DNA sequences obtained from a population*.

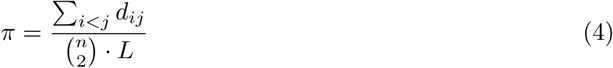

*n is the number of sequences, d_ij_ is the number of different bases between i-th sequence and the j-th sequence. L is the total number of effective loci*.

By analogy to alleles at genomic loci in sequences, distinct values in the 12-dimensional natural vector correspond to genetic variation. Crucially, each vector component represents a continuous magnitude with biological significance. To preserve this quantitative information, the conventional definition of nucleotide diversity (*π*) requires reformulation. Our modified definition explicitly incorporates pairwise differences in vector components, defined as definition **??**:

##### Definition 3

(Pairwise diversity of natural vectors ***π***: ***π_NV_***)

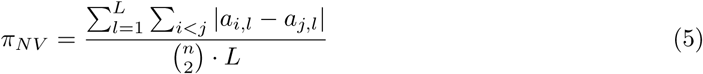

*n is the number of sequences. L is the total number of effective loci. For natural vector, L is the number of dimensions. a_i,l_ is the numerical value of natural vector corresponding to i-th sequence on l-th dimensional site, a_j,l_ is the numerical value of natural vector corresponding to j-th sequence on l-th dimensional site*.

The pairwise diversity of natural vectors in Definition 3 differs from that in Definition 2. Instead of directly counting the number of pairwise differences, it incorporates the magnitude of the absolute differences between the vectors. This is because, in the context of natural vectors, the magnitude of the difference in each dimension carries meaningful biological or statistical implications—such as variations in the size of the second moment. These numerical discrepancies contribute meaningfully to the definition of pairwise diversity of natural vectors.

#### 2.6.3 Shannon’s Entropy

Shannon’s entropy is one method based on the information theory to measure genetic diversity. The calculation formula is as follows:

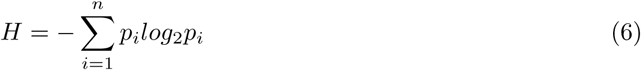

*p_i_*is the frequency of the i-th allele and *n* is the number of alleles. High value of Shannon’s entropy represents high genetic diveristy.

While both Expected Heterozygosity and Shannon’s Entropy are used to measure genetic diversity, they differ in their calculation methods and interpretations. Expected Heterozygosity is more focused on the probability of heterozygosity and is less sensitive to sample size.

### 2.7 Simulation for the mutation effect on NV

Starting from the mitochondrial variation data set of 1000 genomes, the SNP variation data set was sorted out. On the human mitochondrial reference genome, a simulated single-SNP mitochondrial variant sequence was obtained by changing the SNP variants one at a time. For each simulated single-SNPmitochondrial variant sequence, its 12-dimensional natural vector was calculated and compared with the 12-dimensional natural vector calculated for the mitochondrial reference genome. The distance between the two natural vectors was defined by the Euclidean distance between the reference natural vector and the single variant natural vector, that is, the values of all dimensions were difference, squared, summed, and finally squared again. This value is the effect of a single SNP on the natural vector of mitochondrial genome sequence. Similarly, the differences can be calculated as the effect of individual SNP variants on the mean dimension and the effect on the variance dimension for each of the four mean and four variance dimensions, respectively.

### 2.8 Data Set

The 1000 Genomes Project[8] is an international collaborative research initiative aimed at mapping the most detailed and medically valuable human genetic polymorphism map through high-throughput sequencing technology. The reference data resources generated by the project remain heavily used by the biomedical science community. In our research, all modern human mtDNA genomes are downloaded from phase 3 of the 1000 Genomes Project, revised mtDNA calls for 1000 Genomes Phase 3. Total 2534 revised modern human mtDNAs are from 26 populations and cover samples from Africa, Europe, Asia, America and Oceania. Detailed information is listed in the supplemental files.

## 3 RESULTS

### 3.1 Identification of discrete numerical segments of Nautral Vector

Taking the mtDNA Genomes data in 1000Genomes Project as an example, to identify the generalized alleles for each dimension of NV, we aim to distinguish human populations according to their geographical origins: Africa, Eurasia and East Asia. Randomly select a population from each ethnic group and collect mitochondrial data from 20 individuals within that population. We used the British, the Yoruba, and the Han Chinese populations firstly. Subsequently, we applied the discretization algorithm and presented the results in Figure 1 A. Experiments were conducted independently for each dimension. From the figure, most tests could obtain the minimum p-values when the numbers of sub-intervals are less than 10 except for *NV*_8_(*µ_T_*) and *NV*_10_(*D_C_*), however, their p-values are also smaller than 0.05(0.0062 and 0.0033 for each locus) when the number of intervals are chosen as 5. This experiment thus suggests that a detailed segmentation of intervals is unnecessary when applying the discretization algorithm to this particular dataset, a division within 10 sub-intervals is sufficient for the vast majority of cases.

**Figure 1:**
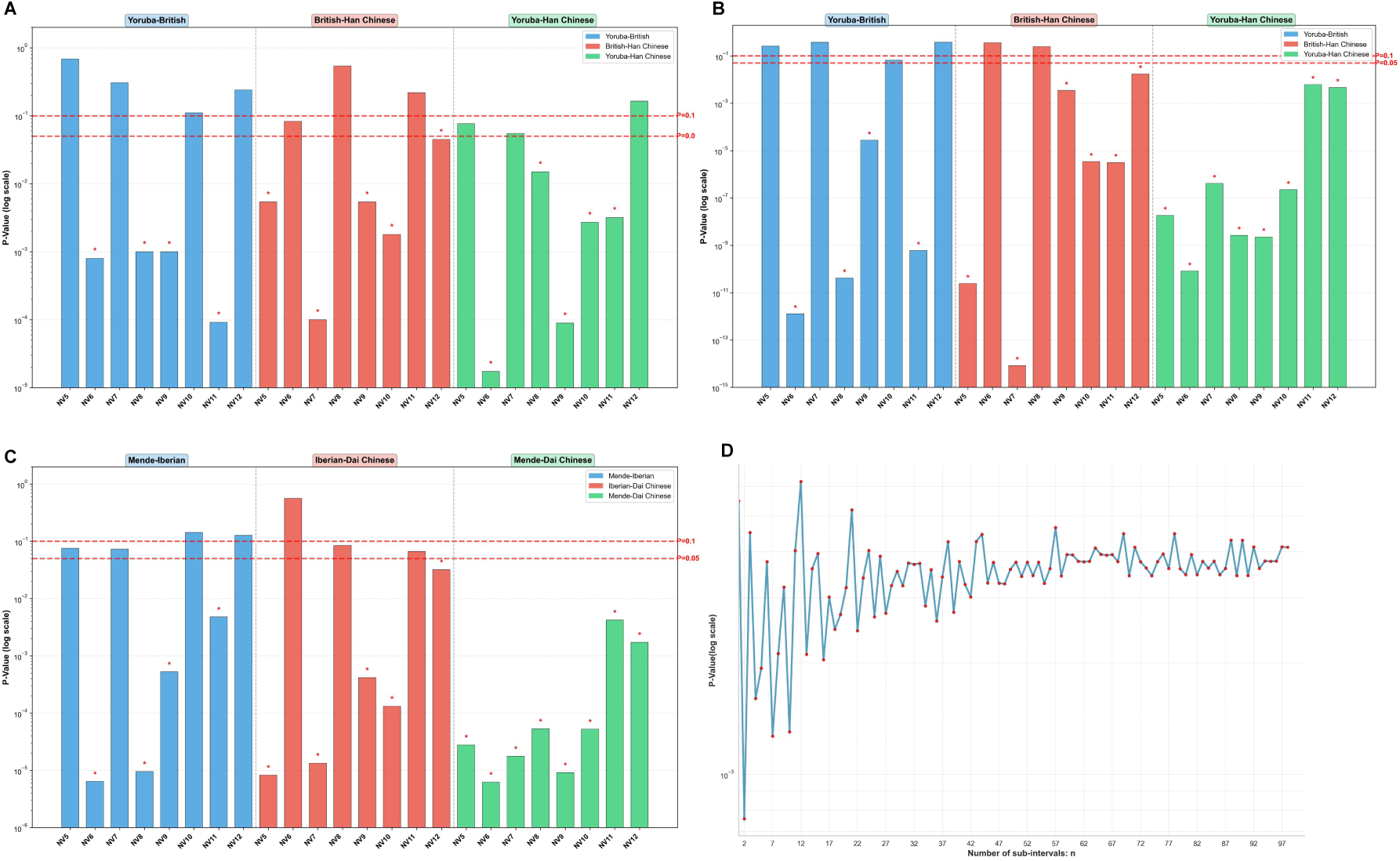
Identification of discrete numerical segments of Nautral Vector. **(A)** Histogram of P-values (on log scale) from paired Wilcoxon tests for three population pairs: Yoruba-British, British-Han Chinese, and Yoruba-Han Chinese, using 60 samples (20 each from Yoruba, British, and Han Chinese). The samples were equally divided to achieve the minimum p-values. The three pairs are represented by distinct colors. Red dashed lines indicate p-value thresholds of 0.05 and 0.1. Bars with p-values below 0.05 are annotated with red stars. **(B)** Similar to (A), but with 303 samples (108 from Yoruba, 92 from British, and 103 from Han Chinese). **(C)** Similar to (A), but for paired Wilcoxon tests involving population pairs: Mende-Iberian, Iberian-Dai Chinese, and Mende-Dai Chinese, using 60 samples. **(D)**P-values as a function of the number of intervals (ranging from 2 to 100) for the Yoruba-British comparison based on the *NV*_6_(*µ_C_*) statistic.

Besides, we conducted an experiment on the detailed effect of changing the number of subintervals on p-values. Through the line chart in Figure 1 D, p-values tend to be small when the number of sub-intervals is smaller than 25 and tend to be stable for large number of sub-intervals. For Yoruba-British’s *NV*_6_(*µ_C_*), the p-value attains the minimum 0.0008 when n=4 and is 0.0045 when n=5, the latter is also a convincing result to deduce population difference. For Yoruba-British’s *NV*_5_(*µ_A_*) shown in supplementary document, p-values are large integrally, nonetheless, the smaller values distribute in the left segment of the line chart. The p-value attains the minimum 0.6872 when n=3 and is 0.8057 when n=5, whichever indicate non-significance for the statistical hypothesis.

A fundamental prerequisite for identifying generalized alleles lies in the premise that populations distinguished by these alleles originate from distinct subpopulations exhibiting well-defined population structures. In the presented example, the three human populations are categorized based on divergent skin pigmentation phenotypes. A critical concerns arise regarding the efficacy of generalized allele identification when either of the following conditions occurs: (1) the target species lacks explicit population population structure because of migration or gene flow, or (2) the populations of the targeted species show close phylogenetic relationship. To test the boundary of the method effectivity, we systematically evaluated this scenario through comparative analysis of two sets of genetically adjacent populations. The first is three populations from east Asia, Han Chinese, Southern Han Chinese, and Japanese. The second set includes three populations from Indo-Aryan branch spoken primarily in South Asia: Bengali, Gujarati and Punjabi. These results of identification are in the 3.2.3 section.

#### 3.1.1 Influence of Population Pick-up to the “Allele” Identification

In the former experiments, a set of samples were chosen from three populations Yoruba, British and Han Chinese and corresponding analyses were based on them. The original intension why these three are firstly selected is they cover three types of modern human, and their combination is one of potential ones after all. Another question worth exploring is whether changing the population while still maintaining the same framework for African, Eurasian and East Asian will affect the stability of our discretization method. Several experiments are designed through changing one or two or three populations. The populations are marked as ‘African population’, ‘Eurasian population’ or ‘East Asian population’ respectively. Populations belonging to ‘African population’ include Gambian, Luhya, Mende and Yoruba; belonging to ‘Eurasian population’ include Finnish, British and Iberian; belonging to ‘East Asian population’ include Japanese, Han Chinese and Dai Chinese.

We designed two groups of experiments: the population combinations in the first group is shown in Table 1 and the other is shown in Table 2. Both groups were applied the controlled variable method for each group, which means one population was altered at a time while keeping the other two populations constant.

**Table 1:**
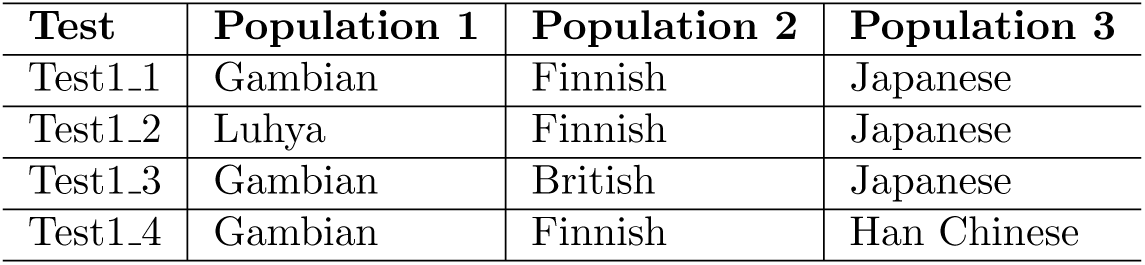
Populations involved in the first group of experiments: four experiments are labeled as Test1 1, Test1 2, Test1 3, Test1 4

| Test | Population 1 | Population 2 | Population 3 |
| --- | --- | --- | --- |
| Test1.1 | Gambian | Finnish | Japanese |
| Test1.2 | Luhya | Finnish | Japanese |
| Test1.3 | Gambian | British | Japanese |
| Test1.4 | Gambian | Finnish | Han Chinese |

**Table 2:** Populations involved in the second group of experiments: four experiments are labeled as Test2 1, Test2 2, Test2 3, Test2 4

| **Test** | **Population 1** | **Population 2** | **Population 3** |
| --- | --- | --- | --- |
| Test2_1 | Mende | Iberian | Dai Chinese |
| Test2_2 | Mende | Iberian | Han Chinese |
| Test2_3 | Mende | British | Dai Chinese |
| Test2_4 | Yoruba | Iberian | Dai Chinese |

While altering one population, most of the minimum p-values can remain within the same order of magnitude. Here, one result of p-values with ‘Mende-Iberian-Dai Chinese’ population combinations is shown in sub-graph C of Figure 1. It can be observed that the results are stable compared to Yoruba, British, and Han Chinese shown in sub-graph A. Besides, results of other population combinations are shown in supplementary document. In sub-graph A, except for ‘Eurasian-East Asian’ pair in Test1 group whose p-value changed from greater than 0.05 to less than 0.05 when Test1 4 was carried out(Finnish-Han Chinese pair), p-values of other pairs kept stable while population combinations changed. For Test2 group, p-values of three pairs kept stable. In sub-graph B, similar results could be observed: p-value of Test1 group of ‘Eurasian-East Asian’ pair changed from less than 0.05 to greater than 0.05 when Test1 4 was carried out, p-values of other pairs kept stable while population combinations changed. And p-values of three pairs kept stable for Test2 group.

Overall, altering one or several populations, but if the three populations are derived from Africa, Eurasia and East Asia respectively, the discretization method we designed is effective and stable.

#### 3.1.2 Influence of Population Size to the “Allele” Identification

In the previous experiments, 60 samples are chosen randomly from sets consisted by three populations: British, Yoruba and Han Chinese; Han Chinese, Southern Han Chinese, and Japanese; Bengali, Gujarati and Punjabi. One question is what would happen if more samples are added to these three populations. In other words, we want to know whether the number of sub-intervals chosen for discretization would be stable. In fact, there are 108 sequences from Yoruba, 92 from British and 103 from Han Chinese in the 2534 sequences. The same experiment was conducted on the 303 sequences (comprising 108, 92, and 103 sequences) and the numbers of sub-intervals for each general locus were kept same, and the results are presented in Figure 1 B. Taking 0.05 and 0.1 as two thresholds of p-value(*p <* 0.05 indicates a significant hypothesis test), changes in p-values between 60 samples and 303 samples can be observed by the changes in color of the two subgraphs after grouping. For each population pair, numerical values in the left columns are p-values and the right columns contain the corresponding number of intervals which are equally divided. Comparing graph A and B, 6 p-values varied in groups, 5 of them varied from group with higher p-values to group with lower values, for example, p-value of NV10 of Yoruba-British varied from 0.1104 to 0.067 when population size increased, which means more samples tend to make test more significant. Other p-values remain within the same order of magnitude. Therefore our discretization algorithm is stable in terms of varying sample numbers to a certain extent.

#### 3.1.3 Influence of Close-related Populations to the “Allele” Identification

Here we identified “allele” for two sets including close-related populations. The first set includes Han Chinese, Southern Han Chinese and Japanese. Japanese genetics show some similarity to both Han Chinese and Southern Han Chinese. And there is a strong genetic similarity between the Han Chinese and Southern Han Chinese. The second set includes Bengali, Gujarati and Punjabi which are all from India and the Indo - Aryan language branch. For east set, 20 individuals were randomly sampled from each of the three populations. The results of “allele” identification are shown in Fig. S6 of supplemental documents.

We still independently discretized each dimension (”locus”) to get the number of sub-intervals, which gives the number of generalized alleles. The results show that since our generalized allele discretization algorithm is based on the division of different populations, when applied to closely related groups, a larger P value will occur, which means that it is difficult to obtain the best number of sub-intervals(generalized alleles) through the minimum P value. So it implys that the identification of “allele” needs clear population structure. It can be observed from the Fig. S6 that in these two sets of data, the minimum p-values of paired wilcox test on 60 samples are mostly greater than 0.1, while the majority of the p-values in the previous results based on the division of African, Eurasian and East Asian populations are less than 0.05. The same experiments were also conducted on the two datasets with varied individual number, whose results obtained were displayed in the supplementary files. This observation demonstrates that a well-defined population structure at the root of intraspecific phylogenetic trees constitutes a critical prerequisite for methodological development. Crucially, when this foundational requirement is satisfied, closely related populations at the crown of the phylogenetic tree can be well distinguished. The following section show that, taking the population genetic diversity parameters as an example, the Pi value (the average number of nucleotide mismatch) of global populations in human species calculated by this method is well fitted with the values calculated by calssical SNP data.

### 3.2 Principal Component Analysis Using Natural Vector and Comparison to SNP-based Results

Principal Component Analysis (PCA) is a widely used statistical method that enables the visualization and quantification of population stratification by reducing high-dimensional genotype data into a few principal components that capture the major axes of genetic variation. In this study, we applied PCA to explore the genetic structure of different populations from 1000Genomes Project and to identify potential clusters or patterns of populations. PCA of populations based on SNP data and natural vectors was performed as described in the Method section. The results are shown in Figure 3.

**Figure 2:**
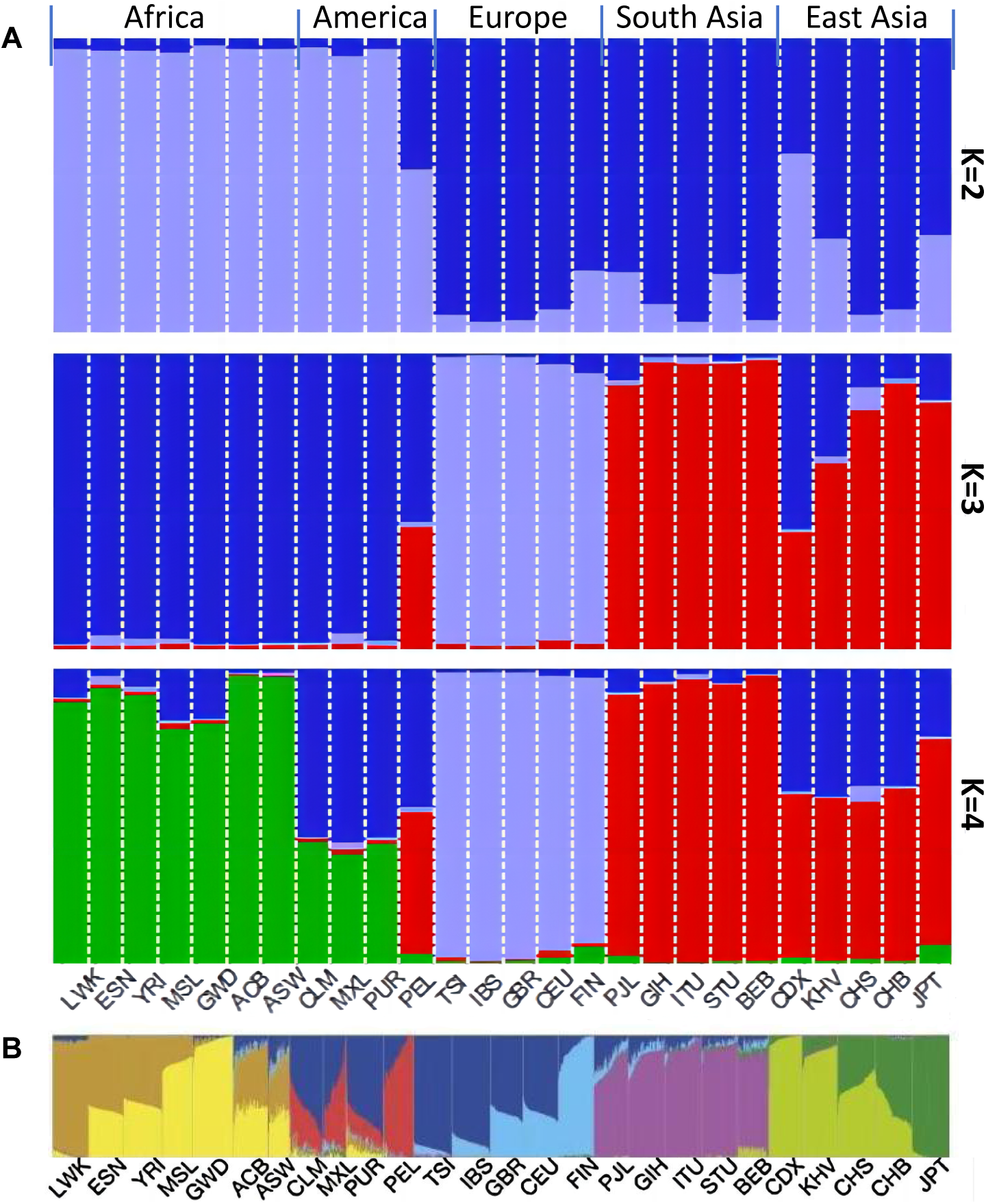
Population Genetic Structure Inferred from Mean Natural Vectors. **(A)** Bar plot generated by plotQ showing the estimated population structure for K=2, 3, and 4 clusters. Each vertical bar represents one of the 26 populations (abbreviations on the x-axis), grouped by their geographic origin: Africa, America, Europe, South Asia, and East Asia. The colors within each bar represent the estimated ancestry proportions (Q-matrix) belonging to the K genetic clusters. The plot demonstrates the hierarchical population stratification at different levels of genetic resolution.**(B)** Individual-level population structure plot for the same 26 populations. Each vertical bar represents a single individual, grouped by their population assignment. This view reveals the distribution of ancestry components within each population, highlighting the degree of admixture or genetic homogeneity at the individual level. The consistent color scheme for ancestry components across both (A) and (B) allows for direct comparison between the population-averaged and individual-specific patterns.

**Figure 3:**
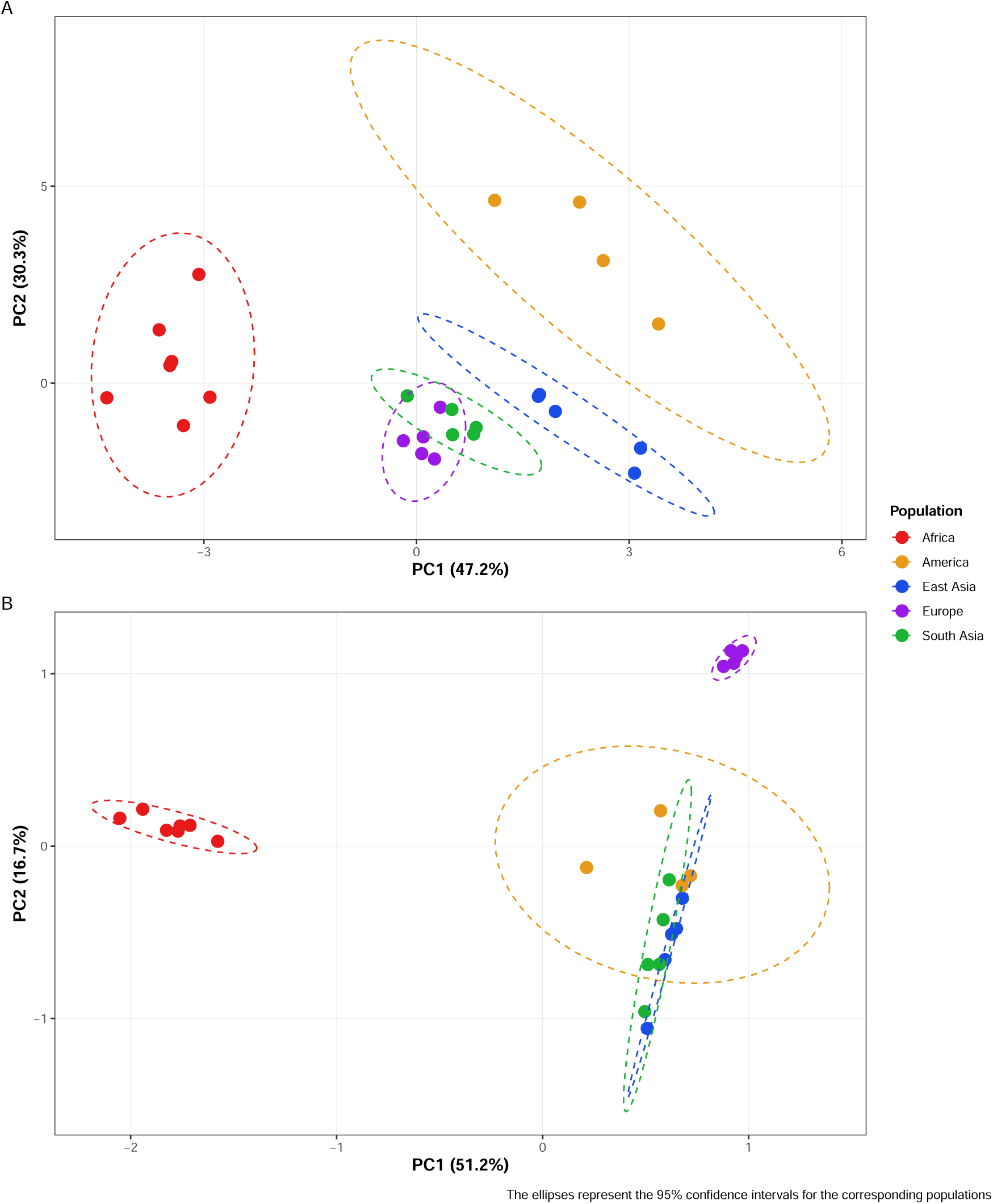
Principal Component Analysis (PCA) of human population clustering based on SNP data and natural vectors. (A) PCA plot of natural vectors (PC1: 47.2% variance explained; PC2: 30.3% variance explained). (B) PCA plot of SNP data (PC1: 51.2% variance explained; PC2: 16.7% variance explained).Points represent populations, colored by their geographic ancestry area (Africa, America, East Asia, Europe, South Asia). Ellipses depict the 95% confidence intervals for each population group.

PCA based on natural vectors revealed distinct clustering patterns across five major global areas (Africa, America, East Asia, Europe, and South Asia). As shown in Figure 3.A, the first two principal components (PC1: 47.2%; PC2: 30.3%) captured 77.5% of the total genetic variation. Populations formed geographically defined clusters, with 95% confidence ellipses showing clear separation between continental groups. Besides Europe and South Asia, populations from other three areas are each gathered together. Points representing populations from Europe and South Asia are closer and their two 95% confidence ellipses overlap.

Comparative analysis with SNP-based PCA (Figure 3.B) demonstrated concordant population stratification. Both methods identified Africa as the most divergent cluster (extreme left in PC1). South Asian groups occupied intermediate positions in both analyses, consistent with known admixture histories. The ellipsoidal dispersion patterns indicate that natural vectors preserve interpopulation relationships while compressing high-dimensional sequence data efficiently as SNP-based approaches. Besides, the cumulative variance explained by the first two principal components (PC1 and PC2) was higher in the analysis based on natural vectors (77.5%) than in the analysis based on SNP data (51.2%).

### 3.3 Genetic Diversity Analysis Using Natural Vector and Comparison to SNP-based Results

Genetic diversity refers to the degree of variation in genetic information within a population and is an essential component of biodiversity. Research on genetic diversity is of great significance for studying the adaptability and evolution of species. Genetic diversity can be calculated and measured through various methods and indicators and results calculated in this work could reflect some features of modern humans from 1000Genomes Project. Detailed results of genetic diversity represented by Pi value from different populations are shown in Figure 4. There are two main phenomenons we can observe from the results.

**Figure 4:**
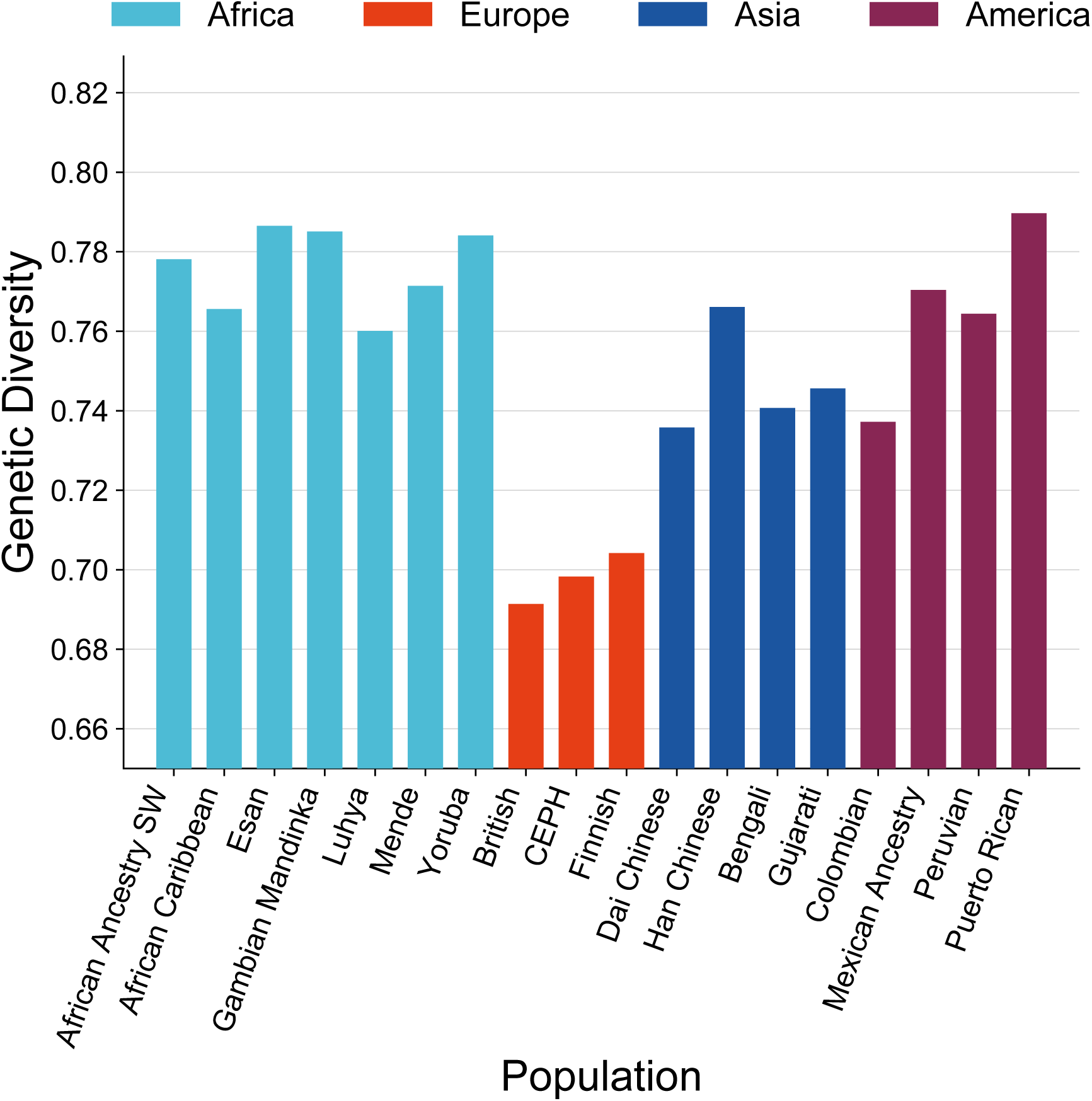
Genetic diversity from mtDNA genomes in 1000Genomes Project. GD is short for genetic diversity

#### 3.3.1 Highest Genetic Diversity of African Populations

From Figure 4, besides populations from America, African populations exhibit the highest genetic diversity among all human populations. This phenomenons can be explained through various genetic markers, such as single nucleotide polymorphisms (SNPs), copy number variations (CNVs), and other forms of genetic variation of African populations. For instance, studies have shown that African populations harbor a significant proportion of the world’s genetic diversity[9]. Genome-Wide Association Studies (GWAS) studies have consistently shown that African populations harbor a large number of unique genetic variants. Whole-Genome Sequencing: Recent whole-genome sequencing efforts have revealed millions of genetic variants in African populations, many of which are rare or unique to Africa. Meanwhile, certain African populations, such as the San people of southern Africa, have been identified as having exceptionally high levels of genetic diversity.

#### 3.3.2 High Diversity in American Related with Gene Flow

Three populations of America: Mexican Ancestry, Peruvian and Puerto Rican exhibit high genetic diversity similar to that of African populations. This phenomenon can be explained through their geographical locations and historical migration. For Mexican, Mexico has a complex and unique history, with its population composed of multiple different ancestral groups[10]. The genome of Mexicans has several different ancestral sources, including indigenous peoples, Europeans, and Africans. This complex historical background has led to a high level of genetic diversity in the Mexican population. For Peruvian, Peru has experienced multiple waves of immigration throughout its history, including Spanish colonizers, African slaves, and immigrants from other regions such as Europe and Asia[11]. Besides, Puerto Ricans are an ethnic group primarily composed of various mixed race individuals, including Indo European, Indo African, and African Europen mixed races.

In summary, the high levels of genetic diversity in the populations of Mexico, Peru, and Puerto Rico are mainly due to historical migration and hybridization, geographic isolation and gene flow[12]. These factors working together have led to the complexity of genetic diversity among populations in these regions. Detailed values of genetic diversity calculations are listed in the supplementary file.

#### 3.3.3 Correlation of Genetic Diversity Under SNP and Natural Vector

When estimating genetic diversity across populations, two distinct estimators of nucleotide diversity (*π*) were employed: the conventional SNP-based *π* derived from genotype matrices and our natural vector-derived *π* (denoted *π_NV_*).

As shown in Figure 5.A, linear regression analyses revealed strong concordance between *π* of SNP and *π_NV_* across main areas (Africa, Asia, America and Europe), which confirms that our natural vector formulation preserves fundamental diversity signals captured by traditional indices. Among the diversity values, African populations exhibited the highest diversity values, followed by Asian and American groups, while European samples showed the lowest diversity. One phenomenon can be obeserved from plot of *π_NV_* versus *π* (Figure 5 B), is that the data points, while showing a strong positive correlation, are more spread out along the *π_NV_* axis. This wider dispersion of values according to population origin underscores the greater sensitivity of *π_NV_*in capturing inter-population genetic differences.

**Figure 5:**
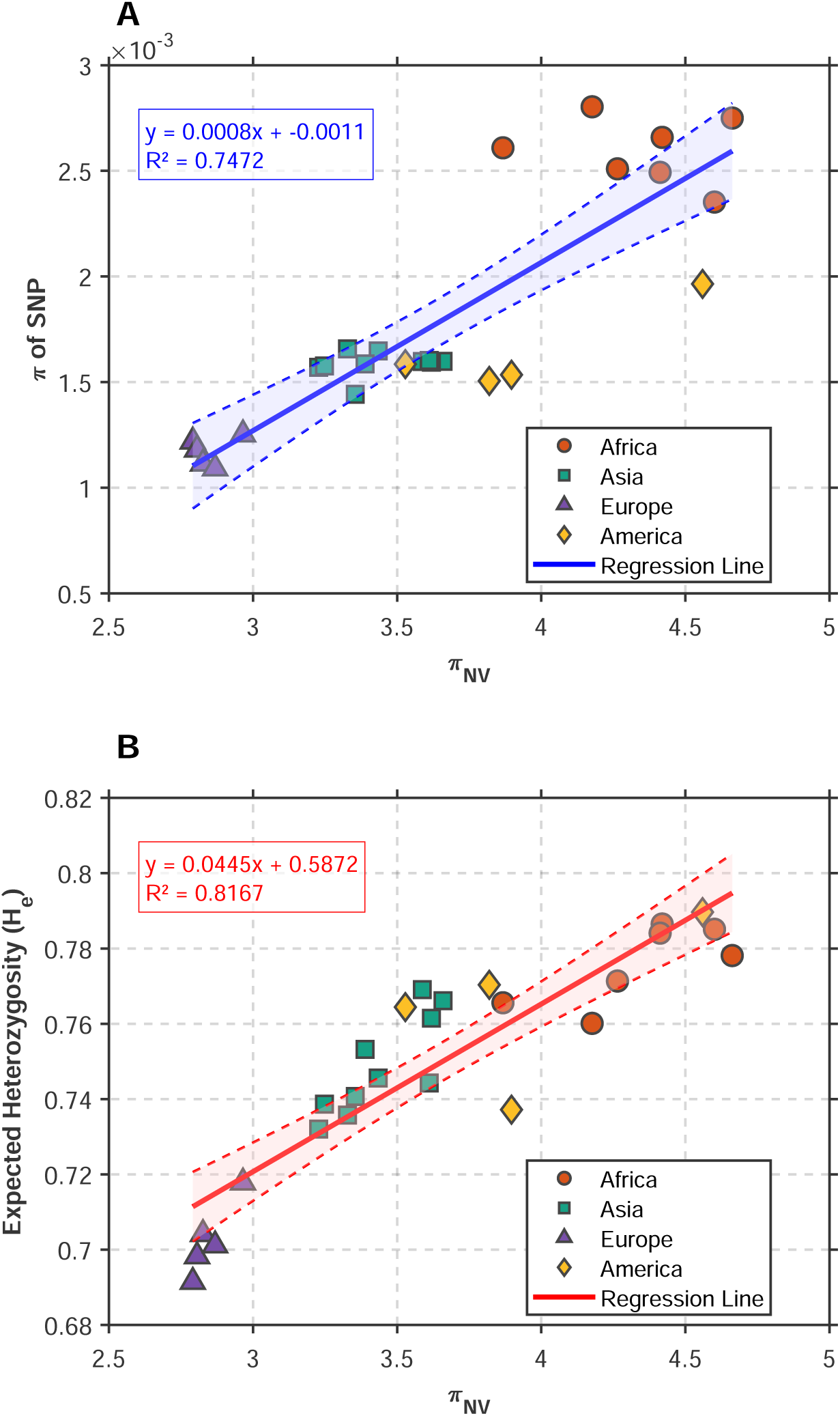
Correlation Analysis of Genetic Diversity Indice (A) Linear regression between traditional nucleotide diversity (*π*) and *π_NV_* (R² = 0.747). (B)Linear regression between expected heterozygosity (*H_e_*) and nucleotide diversity derived from natural vectors (*π_NV_*) across four continental populations (R² = 0.817).Each point represents a population group. The solid line denotes the regression line, with the corresponding equation and coefficient of determination (R²) displayed on the panel.

Besides, as another measurement of genetic diversity, Expected Heterozygosity (*H_e_*) demonstrated a strong positive correlation with newly defined diversity *π_NV_* as shown in Figure 5.B. This demonstrates the robustness of natural vectors in computing genetic diversity.

### 3.4 Mutation of DNA effects on Natural vector

As shown in the definition of the natural vector, the position of a single SNP variant in the sequence directly affects the values across different dimensions of the natural vector. Simulation results confirm this conjecture.

According to Figure 6 A and B, the effect of point mutations is mainly influenced by the location where the mutation occurs. As for the influence of mean dimenstions in Figure 6 A the closer to the middle of the sequence, the smaller the effect, and the closer to the ends of the sequence, the greater the effect. The effect of proximity to the 5’ or 3 ‘end of the sequence was symmetric. The same mutation type would show a consistent linear pattern of mutations. Figure 6B showed the effect of point mutations on the four variance dimensions. It can be seen that the variance dimension was affected by the location of the mutation, which was a complex “bird” shape of pattern, but there was also consistency in symmetry and mutation type patterns.

**Figure 6:**
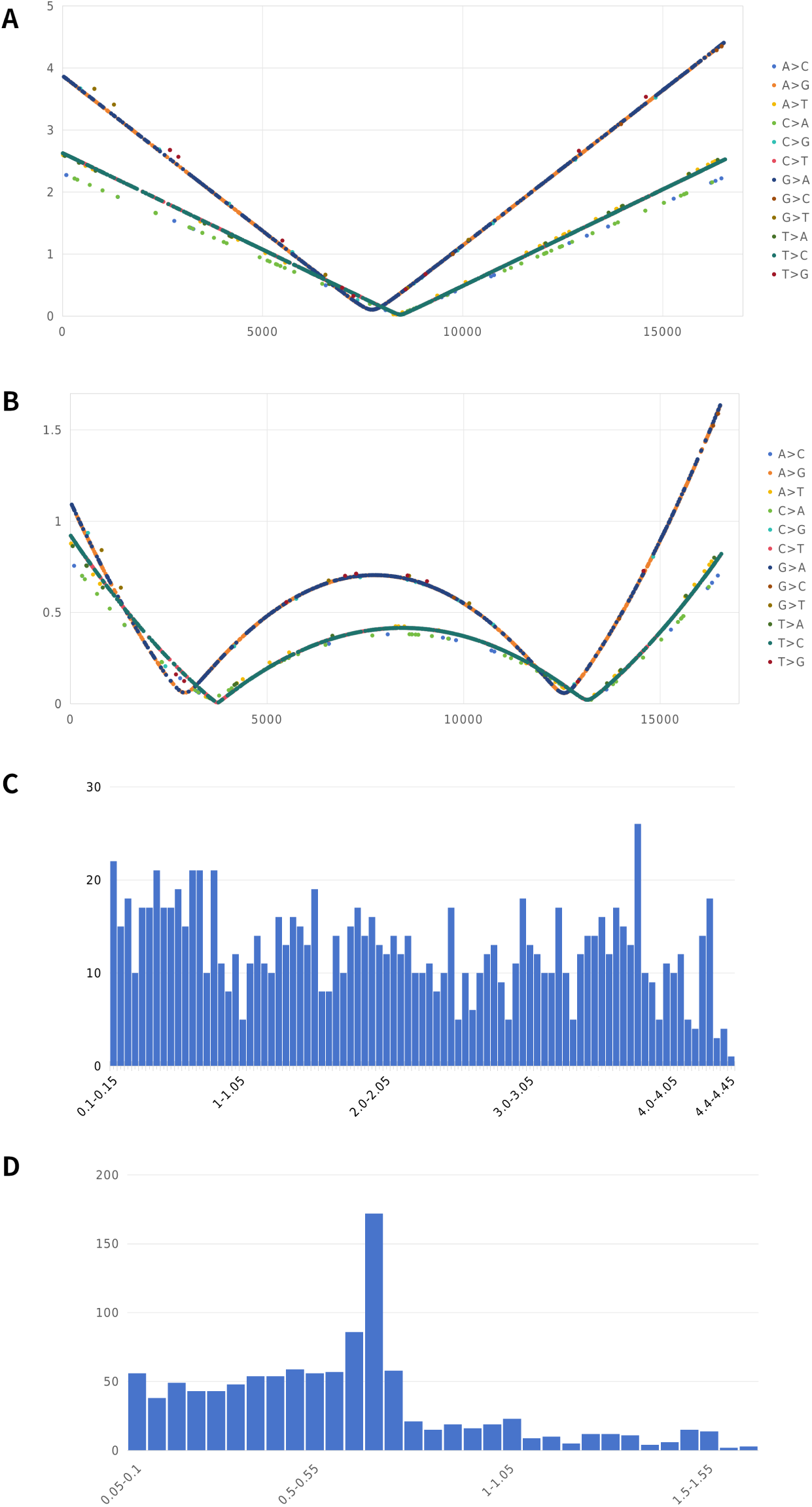
(A) The positional effect of a single SNP variant on the natural vector values. (B) The positional effect of the same variant type on different dimensions of the natural vector. (C) The overall distribution of the effect magnitude for all single SNP variants on the natural vector, regardless of position or type.(D) The distribution of the effect magnitude for a single type of SNP variant.

Additionally, two notable phenomena deserve attention. Firstly, different types of single SNP variants exhibit distinct effects on the natural vector. Moreover, this concave trend appears to be divided into several different ‘concave curves’. The two most frequent types of mutations, A*>*G and C*>*T, form the most “densely packed” curves. Although other mutation types are relatively rare, it is evident that mutations of the same type do not appear on different “concave curves”.

When examining different types of SNP variants separately, it can be observed that although the number of variants varies significantly across types, variants of the same type entirely lie on the same “concave curve”. This suggests that different types of variants exhibit distinct positional effects. Second, even for the same type of variant, the effects on different dimensions of the natural vector can vary considerably. As shown in Figure 6A, for dimensions representing the mean positions of the four nucleotides, the effect of a variant shows a strong linear relationship with its position. In contrast in Figure 6B, for dimensions representing the variance of the four nucleotides, the effect exhibits a unique double-concave trend.

If the positional effect is disregarded, the effect of a single SNP variant on the natural vector values can be treated as a random variable. Figure 6C and D illustrated the distribution of mutation effects on position and on effect size size using the A*>*C variant as an example. In particular, it can be seen from Figure 6D that there was a sharp peak in the effect of variants of the same type on the natural vector.

## 4 Discussion

We applied the natural vector method to the population genetic analysis of a sequence sample. We treated the natural vector method as a kind of abstract “molecular sequence markers” and analogized it with genetic markers such as SNP. Namely, we analogized dimensions as some kinds of gerneralized “genetic loci” and dimensional numbers as “generalized alleles”, so that multiple population genetic analysis methods can be transferred to the natural vector method. A very important issue here was how numerically continuous dimensions in natural vectors can be discretization to produce population frequencies and satisfy the requirement that the true allele is finite. We solved this problem by establishing an optimal discretization algorithm based on the degree of differentiation of the meta-population. The results obtained are quite similar to those of SNP-based population genetic analysis. In some aspects, such as the ability to cover more different types of variation, natural vectors are more advantageous than SNP-based approach.

The natural vector method is based on that the whole DNA sequence is known, but population genomics analysis is based on the identification of variation. Thus, for most multicellular animal and plant species, low-coverage DNA sequencing data has provided good enough variant-calling power but are not sufficient to generate a complete DNA sequence assembly. Therefore, majority of the existing population genome data cannot be directly used for natural vector analysis. This limitation of known whole DNA sequences is an important reason why we chose mitochondrial genome data instead of nuclear genome data in this study. This limitation also makes haplotyping of sexually reproducing diploid and even polyploid species an important difficulty for the application of natural vector methods. The analysis of diploid heterozygote and even haplotype is an important difficulty in population genetic analysis using natural vector method in the future.

The problem of representing mutation has long been recognized as a difficulty in the application of alignemnt-free methods to population genetic analysis [13]. In this paper, we present an analysis of the effects of a single point mutation of SNP on different dimensions of natural vectors. In fact, this idea is inspired by the Fisher geometric model [14], which is to manipulate the geometric space of natural vectors by analogy to the phenotypic space. We can see that the “effect” of a mutation on a natural vector (which can be seen as a kind of analogy to the phenotypic effect of a mutation) depends on the position of the mutation in the sequence and the type of mutation. At the same time, the effect distribution of mutations may follow some common unimodal distribution controlled by the parameters. This is an essential starting point on the investigation of muation and alignment-free methods in population genetic analyses. In the future, it is necessary to further explore the relationship between mutations and natural vectors within the framework of genetics, involving issues includes whether the effect of mutations on natural vectors is related to fitness, the relationship between the frequency of mutations and the effect of natural vectors, and the relationship between the distribution of mutations on the sequence and the effect of natural vectors.

## List of Abbreviations

GD: Genetic Diversity
mtDNA: Mitochondrial DNA
NV: Natural Vector

## 5 Declarations

### 5.1 Ethics approval and consent to participate

Not applicable.

### 5.2 Consent for publication

All the authors approved the publication of the manuscript.

### 5.3 Availability of data and materials

The data used in this study are publicly available and can be accessed through the phase 3 of the 1000 Genomes Project, revised mtDNA calls for 1000 Genomes Phase 3. Also, interested researchers can request access to the data by contacting the corresponding author or other authors.The authors affirm that all data necessary for confirming the conclusions of the article are present within the article, figures, and tables.

### 5.6 Authors’ contribution

Project design and conceptualization was performed by Stephen S.-T.; Data curation, Formal analysis, Investigation, and Writing - original draft were performed by Mengcen Guan and Qi WU; Writing - review & editing were performed by Mengcen Guan, Qi WU and Xin Zhao. All authors have read and agreed to the published version of the manuscript.

## 5.7 Acknowledgments

Not applicable.

